# Progressive suppression of DNA repair genes with persistent p53 activation in Doxorubicin-treated cardiomyocytes

**DOI:** 10.64898/2026.02.03.703628

**Authors:** Emma M. Pfortmiller, José A. Gutiérrez, Alyssa R. Bogar, Michelle C. Ward

## Abstract

Doxorubicin (DOX) is an effective anti-cancer drug; however it can cause cardiotoxicity by inducing DNA double-strand breaks in cardiomyocytes. Cardiotoxicity can manifest immediately or years following treatment. Most human *in vitro* models of DOX-induced cardiotoxicity (DIC) focus on the acute effects of DOX treatment. To understand the long-term effects, we profiled the global gene expression response to DOX exposure over time. We treated iPSC-derived cardiomyocytes from six individuals with DOX for 24 hours and assayed responses after 0, 24 and 144 hours of recovery. DNA damage, determined by γH2AX expression, is induced following DOX treatment and is resolved by the final recovery timepoint. We identified both acute and chronic gene expression response signatures. The chronic signature, representing 501 genes, is enriched for p53 target genes and DNA damage response genes compared to acute response genes. P53 target genes are persistently activated, and DNA damage response genes are progressively downregulated over time. Our results suggest an altered cell state following repair of double-strand breaks that is distinct from pre-exposed cells. DOX response genes with persistent changes in expression can be applied to the design of toxicity biomarkers or therapeutic targets.

## Introduction

Doxorubicin-induced cardiotoxicity (DIC) is a well-established clinical phenomenon that affects a subset of cancer patients treated with the drug (Camilli *et al*. 2024). Cardiotoxicity can lead to cardiac dysfunction and ultimately heart failure (Cardinale *et al*. 2020). DIC can occur soon after or years following treatment. Typically, cardiotoxicity is diagnosed once damage has occurred. For example, elevated levels of cardiac troponin I in the blood indicate that the heart is under stress. Cardioprotective agents can be administered to prevent cardiotoxicity; however, there is currently only one FDA-approved drug, dexrazoxane, that is in limited use given its reduced anti-cancer efficacy. There is therefore a need to identify both early biomarkers of toxicity and potential therapeutic targets.

Doxorubicin (DOX) is a Topoisomerase II inhibitor (TOP2i) that induces DNA double-strand breaks. The molecular basis of DIC is challenging to study in humans *in vivo* given the inaccessibility of heart muscle. Induced pluripotent stem cell-derived cardiomyocytes (iPSC-CMs) have been shown to effectively model the disease (Burridge *et al*. 2016). Treating iPSC-CMs with sub-micromolar concentrations of DOX induces large scale changes to the transcriptome (Knowles *et al*. 2018; Matthews *et al*. 2024) and the proteome (Johnson *et al*. 2025). iPSC-CMs have also been used to identify mediators of DOX-toxicity using functional genomic CRISPR screens, which have highlighted the involvement of transporters such as SLC01A2 and metabolic enzymes such as carbonic anhydrase (Sapp *et al*. 2021; Liu *et al*. 2024). CRISPR knockdown of genes in genetic loci associated with DIC have also highlighted genes most likely to be associated with the disease (Fonoudi *et al*. 2024). All the studies described above measure the effects of DOX within 72 hours of treatment.

Clinically, DOX is typically administered through week-long cycles which include recovery periods between treatments. Human autopsies of individuals treated with DOX ante-mortem have detectable levels of DOX in their heart tissue suggesting that long-term exposure of cardiomyocytes may have long-term effects (Stewart *et al*. 1993). Chronic DOX treatments have been administered in mouse and rat *in vivo* models of cardiotoxicity (Zhu *et al*. 2008; Podyacheva *et al*. 2021); however these studies are unable to detect human-specific responses. A repeated exposure framework has been implemented in studies using iPSC-CMs from a single individual which have identified effects on miRNA expression, metabolites and cardiomyocyte contraction indicating likely dynamics in treatment responses (Chaudhari *et al*. 2016a; Chaudhari *et al*. 2016b; Chaudhari *et al*. 2017).

To capture a chronic, rather than acute DOX exposure state in human cardiomyocytes that can contribute to the development of biomarkers, we designed a study to identify gene expression changes that persist in iPSC-CMs following DOX exposure. We treated iPSC-CMs from six individuals with a sub-lethal DOX concentration for 24 hours and measured the effects 0, 24 and 144 hours following the removal of the drug. This allowed us to identify both acute and chronic gene expression patterns and characterize their properties.

## Methods

### Induced pluripotent stem cell lines

iPSCs were sourced from the iPSC Collection for Omic Research (iPSCORE) Resource, generated by Dr. Kelly A. Frazer at the University of California San Diego as part of the National Heart, Lung, and Blood Institute Next-Gen Consortium (Panopoulos *et al*. 2017). iPSC lines, reprogrammed from skin fibroblasts, may be obtained through contacting Dr. Kelly A. Frazer at the University of California San Diego or through the biorepository at the WiCell Research Institute (Madison, WI, USA).

We utilized six human iPSC lines derived from three males and three females between the ages of 18 and 32 years. These individuals are unrelated and have no prior history of disease. Individual details: Individual 1: UCSD143i-87-1, Female, age 21, Asian (Chinese); Individual 2: UCSD132i-78-1, Female, age 21, Asian (Chinese); Individual 3: UCSD129i-75-1, Female, age 30, Asian (Irani); Individual 4: UCSD138i-84-1, Male, age 21, Asian (Chinese); Individual 5: UCSD178i-17-3, Male, age 18, Asian (Japanese); Individual 6: UCSD154i-90-1, Male, age 23, Asian (Chinese).

### iPSC maintenance

iPSCs were cultured in feeder-free conditions at 37 °C with 5% CO_2_ and atmospheric oxygen. iPSCs were maintained with mTESR1 (85850, Stem Cell Technology, Vancouver, BC, Canada) media with 1% Penicillin/Streptomycin (30-002-Cl, Corning, Bedford, MA, USA) on Matrigel Matrix (354277, Corning, Bedford, MA, USA) at a 1:100 dilution. iPSCs were passaged every 3-5 days as they reached a confluency of 60-80%.

### Cardiomyocyte differentiation

iPSCs were differentiated into cardiomyocytes as previously described (Matthews *et al*. 2024). iPSCs were seeded onto a Matrigel-coated 100 mm cell culture dish and maintained until reaching 85-90% confluency. The initial step to begin differentiation (Day 0) includes adding the GSK3 inhibitor CHIR99021 trihydrochloride (4953, Tocris Bioscience, Bristol, UK) in Cardiomyocyte Differentiation Media (CDM) containing 500 mL RPMI 1640 (15-040-CM, Corning, Bedford, MA, USA), 10 mL B-27 minus insulin (A1895601, ThermoFisher Scientific, Waltham, MA, USA), 5 mL of GlutaMAX (35050-061, ThermoFisher Scientific) and 5 mL of Penicillin/Streptomycin (100X) (30-002-CI, Corning) to yield a final concentration of 12 μM. Media was replaced with CDM after 24 hours (Day 1). After 48 hours (Day 3), 2 μM of the Wnt signaling inhibitor Wnt-C59 (5148, Tocris Bioscience) was added to cells in CDM. CDM was replaced on Days 5, 7, 10, and 12. Cells start spontaneously contracting between Day 7-10. iPSC-CMs were selected using lactate-containing, glucose-free Purification Media (500 mL RPMI without glucose (11879, ThermoFisher Scientific), 106.5 mg L-Ascorbic acid 2-phosphate sesquimagnesium salt (sc228390, Santa Cruz Biotechnology, Santa Cruz, CA, USA), 3.33 mL 75 mg/mL Human Recombinant Albumin (A0237, Sigma-Aldrich, St Louis, MO, USA), 2.5 mL 1 M lactate in 1 M HEPES (L(+)Lactic acid sodium (L7022, Sigma-Aldrich)), and 5 mL Penicillin/Streptomycin (100X) (30-002-CI, Corning). Purification media was replaced on Days 14, 16, and 18. On Day 20 purified iPSC-CMs were dissociated from the culture plate with 0.05% trypsin/0.53 mM EDTA (MT25052CI, Corning) and counted using a Countess 2 machine. iPSC-CMs were plated in a galactose-containing, glucose-free Cardiomyocyte Maintenance Media (CMM) (500 mL DMEM without glucose (A14430-01, ThermoFisher Scientific), 50 mL FBS (MT35015CV, Corning), 990 mg Galactose (G5388, Sigma-Aldrich), 5 mL 100 mM sodium pyruvate (11360-070, ThermoFisher Scientific), 2.5 mL 1 M HEPES (H3375, Sigma Aldrich), 5 mL 100X GlutaMAX (35050-061, ThermoFisher Scientific), and 5 mL Penicillin/Streptomycin (100X)(30-002-CI, Corning) to metabolically mature iPSC-CMs. iPSC-CMs were maintained in culture for an additional 5-10 days. CMM was replaced on Days 23, 25, 27, 29.

### Cardiac troponin T flow cytometry for iPSC-CM purity determination

Following iPSC-CM differentiation, culture purity was assessed using flow cytometry for cardiac troponin T (TNNT2) expression. Day 25 ± 1 iPSC-CMs were released from the cell culture plate using 0.05% trypsin/0.53 mM EDTA (MT25052CI, Corning) and quenched with CMM. Cell suspensions were strained to generate single cells. A total of 3 million cells were divided into one sample well (TNNT2 and live/dead stain) and three control wells (TNNT2-only, live/dead stain-only and unlabeled) in a deep-well U-bottom plate (13-882-234, BrandTech Scientific, Essex, CT, USA). Cells were stained with a viability marker, Zombie Violet Fixable Dye, diluted in PBS (423113, BioLegend, San Diego, CA, USA) for 30 min at 4 °C in the dark. Cells were washed with PBS and then autoMACS running buffer (130-091-221, Miltenyi Biotec, San Diego, CA, USA). Cells were fixed and permeabilized using the FOXP3/Transcription Factor Staining Buffer Set (00-5523, ThermoFisher Scientific) for 30 min at 4 °C in the dark. Cells were washed and stored in autoMACS running buffer O/N. Cells were incubated with 5 μL Cardiac Troponin T Mouse, PE, Clone: 13-11, BD Mouse Monoclonal Antibody (564747, BD Biosciences, San Jose, CA, USA) diluted in Permeabilization Buffer from the FOXP3/Transcription Factor Staining Buffer Set (00-5523, ThermoFisher Scientific) for 40-50 min in the dark at 4 °C. Cells were washed twice with Permeabilization Buffer and resuspended in autoMACS buffer. A BD LSR Fortessa Cell Analyzer was used to analyze ten thousand cells per sample.

The proportion of live cells expressing TNNT2 was calculated as follows: 1. Cellular debris was excluded by adjusting forward and side scatter. 2. Single-celled populations were gated by forward scatter height and area. 3. The Zombie Violet-positive population was excluded as dead cells. 4. TNNT2 positivity in the live population was determined based on the unlabeled cell population.

### Drug stock preparation

Doxorubicin hydrochloride (D1515; Sigma-Aldrich) was dissolved in DMSO to create a 10 mM stock solution. Aliquots of 100 μL were prepared in separate tubes and stored at −80 °C. A working stock of the desired concentration was prepared in CMM immediately preceding treatment. All DMSO vehicle (VEH) control treatments were prepared using identical volumes to the DOX working stock.

### PrestoBlue cell viability assay

Three individuals (Individuals 2, 3 and 5) were utilized to perform cell viability assays on Day 32-35 iPSC-CMs. Six 96-well plates were seeded with 50,000 iPSC-CMs excluding the outer wells. Prior to treatment, a baseline cell viability measurement (time 0) was taken using the PrestoBlue Cell Viability assay (A13262, Invitrogen, Waltham, MA, USA) according to manufacturer instructions with a one-hour incubation period. iPSC-CMs were treated with a range of eight DOX concentrations (0.01 μM – 1000 μM) and volume-matched VEH in quadruplicate for 24 hours. Controls included eight CMM-only iPSC-CM wells per timepoint and six cell-free wells per plate. Treatments were randomized with the Well Plate Maker (wpm) package in R (Borges *et al*. 2021). Viability was determined with the PrestoBlue assay 24 hours following treatment (time 24). Fluorescence was determined using a Biotek Synergy H1 plate reader with an excitation/emission of 460/490 nm.

To determine cell viability at each time point, background fluorescence was first averaged from the six cell-free wells and subtracted from all samples. This yielded relative fluorescence units (RFLU) for each sample. RFLU values from samples on each plate were divided by the CMM-only iPSC-CM wells, and normalized by the average DOX concentration-matched VEH wells. Time 24 data was divided by time 0 data to yield a relative percent cell viability. A dose-response curve for each individual was generated using the drc package in R which calculated LD50 values for each individual at each timepoint (Ritz *et al*. 2015).

### γH2AX immunofluorescence staining and quantification

iPSC-CMs for three Individuals (Individual 1, 3, and 4) were plated at 150,00 cells per well in 24-well plates. To enable comparison of γH2AX staining across time, we maintained the time of staining constant and varied the starting treatment time. First, Day 27 iPSC-CMs were treated with 0.5 μM DOX and VEH in duplicate for 24 hours, then incubated in CMM only for the next 144 hours (tx+144). All remaining wells received CMM only. On Day 29, CMM was added to all wells. On Day 32, four untreated wells were treated with DOX or VEH in duplicate for 24 hours, then recovered for 24 hours in CMM (tx+24). All remaining wells received CMM. On Day 33, four remaining wells were treated with DOX and VEH in duplicate for 24 hours (tx+0). On Day 34, immunofluorescence staining was performed on all treatment timepoints tx+0, tx+24, tx+144, and secondary antibody-only control wells. iPSC-CMs were fixed in 4% paraformaldehyde for 15 min at room temperature. Next, cells were permeabilized for 10 min in 0.25% DPBS-T (0.25% Triton X-100 in DPBS) and washed with cold DPBS. Cells were blocked in 5% BSA in DPBS-T for 30 min at room temperature and incubated overnight at 4 °C with anti-phospho-Histone H2A.X (Ser139) rabbit monoclonal antibody (1:500; NC1602516; Fisher Scientific) prepared in 1% BSA in DPBS-T. Cells were washed with cold DPBS and incubated with a Donkey anti-Rabbit Alexa Fluor 594-conjugated secondary antibody (1:1000; A21207, Invitrogen) for one hour. Nuclei were counterstained with Hoechst 33342 (PI62249; Thermo Scientific) for 10 min in the dark. Fluorescence imaging was performed on an EVOS microscope. Exposure settings were standardized across all conditions.

Quantification of γH2AX positive nuclei was performed using the Cell Counter plugin in Fiji/ImageJ version 1.54i (Schindelin *et al*. 2012). At least 100 nuclei were counted across three fields of view across duplicate wells per condition. γH2AX presence was scored across nuclei. γH2AX Texas Red pixel intensity was measured within nuclei and divided by nucleus area to yield normalized γH2AX intensity per nucleus. Within each image, the mean of γH2AX intensity across all nuclei was calculated. Three images with consistent nuclei counts were selected for each condition. Intensities were normalized across images by dividing by the highest intensity measurement in each image. The mean γH2AX intensity across these three images were calculated to yield Individual-level γH2AX mean intensity. A t-test was performed to compare γH2AX between DOX- and VEH-treated iPSC-CMs across three individuals at each timepoint. *P* < 0.05 is considered significant. To further assess differences in γH2AX expression over time, a t-test was performed between DOX-treated iPSC-CMs at each timepoint. *P* < 0.05 is considered significant.

### iPSC-CM treatment and collection

1.5 million iPSC-CMs were plated per well of a 6-well plate. Day 29-30 iPSC-CMs from six individuals were treated with 0.5 μM DOX and matched VEH in CMM across three six-well plates. The differentiation and treatment was replicated in one of the individuals (Individual 6). Following 24 hours of treatment, one plate of DOX and VEH was harvested (tx+0). CMM was replaced on the remaining plates. Following 24 hours of recovery, one plate of DOX and VEH was harvested (tx+24). CMM was replaced 48, 96 and 144 hours after treatment (Day 32 +/-1, Day 34 +/-1 and Day 36 +/-1). Following 144 hours of recovery, the final DOX- and VEH-treated plate was collected (tx+144). At each timepoint, iPSC-CM cells were harvested on ice, flash-frozen and stored at −80 °C.

### RNA extraction

RNA was extracted from 42 flash-frozen iPSC-CM cell pellets in treatment-balanced batches. All six treatment conditions per individual were extracted in one batch. RNA was extracted across seven batches corresponding to the six individuals and one replicated individual. Total RNA was extracted using the RNeasy Mini Kit 250 (74106; QIAGEN, Germantown, MD, USA) following instructions from the manufacturer. An Agilent 2100 Bioanalyzer was used to evaluate RNA integrity and concentration. All samples exceeded an RNA Integrity Number (RIN) of 8.0.

### RNA-seq library preparation

dT-selected stranded RNA-seq libraries were prepared from 250 ng of total RNA from each sample, with the exception of the six samples from Individual 1 (35-169 ng). Poly(A) RNA was isolated using the NEBNext Poly(A) mRNA Magnetic Isolation Module (E7490L, New England Biolabs, Ipswich, MA, USA). RNA-seq libraries were prepared in a single batch using the NEBNext Ultra II RNA Library Prep with Sample Purification Beads 96 reactions kit (E7775K, New England Biolabs), then indexed using the NEBNext Multiplex Oligos for Illumina (96 Unique Dual Index Primer Pairs, E6440S). Following library preparation, quality was assessed on an Agilent 2100 Bioanalyzer prior to quantification, pooling, and sequencing. All 42 samples were sequenced paired-end 75 bp across two lanes of the Element AVITI (Element Biosciences, San Diego, CA, USA), which yielded a median of 33,462,422 paired-end read fragments across samples.

### RNA-seq analysis

Raw sequencing reads were assessed for quality using FastQC v0.12.1 (Andrews 2010). Reads were aligned to the human hg38 reference genome using Subread (Liao *et al*. 2013), and read numbers across annotated genes quantified using the featureCounts function (Liao *et al*. 2014). We transformed raw counts to log_2_ counts per million (log_2_ cpm) using the edgeR package in R (Robinson *et al*. 2010). Lowly expressed genes with a mean log_2_ cpm < 0 across samples were excluded, resulting in a final set of 14,319 expressed genes.

#### Principal component analysis

Principal component analysis (PCA) was performed on log_2_ cpm values to identify the major contributors to variation in these data. The prcomp function in the stats package v4.4.2 (Team 2024) was used to generate PCA data, and visualized with the autoplot function of ggplot2 v3.5.2 (Wickham 2016).

#### Correction for unwanted variation

Factors contributing to unwanted variation were identified using the RUVSeq package in R (Risso *et al*. 2014). The RUVs function was used to estimate factors contributing to variation using the replicated Individual 6 data. Data segregated based on DOX treatment time following correction with one factor.

#### Differential expression analysis

We randomly selected one of the replicated Individual 6 treatment sets for pairwise differential expression analysis using the edgeR-voom-limma pipeline (Law *et al*. 2018). We used fixed effects for the treatment, a covariate for the RUV factor, and a random effect for individual implemented through the duplicateCorrelation function. DOX samples at each timepoint were contrasted with timepoint-matched VEH samples. Genes with a Benjamini-Hochberg corrected *P* value < 0.05 are considered DEGs.

#### Gene expression trajectory analysis

The Cormotif package was utilized to jointly model expression patterns across time (Wei *et al*. 2015). Cormotif is a Bayesian clustering approach that identifies patterns that best fit the data and defines them as correlation motifs. We used log_2_ cpm of RUVs normalized counts as input and contrasted DOX-VEH pairs across timepoints. Bayesian Information Criterion (BIC) and Akaike Information Criterion (AIC) analysis identified two motifs that best fit the data. We assigned genes to motifs based on their likelihood of belonging to any motif (clustlike) and the posterior probability of belonging to a specific motif (p.post). Genes were assigned to Motif 1 (Acute) with a clustlike probability > 0.8, p.post > 0.05 in tx+0, and p.post < 0.5 in tx+24 and tx+144 timepoints. Genes were assigned to Motif 2 (Chronic) with a clustlike probability > 0.5, p.post > 0.3 in tx+0, p.post > 0.5 in tx+24, and p.post > 0.5 in tx+144. Using these parameters, 56.9% of all DEGs are assigned to one of the two motifs.

#### Gene ontology and pathway enrichment analysis

We performed gene ontology analysis for Acute and Chronic response genes against a background set of all expressed genes. The gProfiler2 tool in R was used to perform gene set enrichment analysis for each gene set (Kolberg *et al*. 2020). Kyoto Encyclopedia of Genes and Genomes (KEGG) Pathways with an adjusted *P* < 0.05 are considered significantly enriched.

#### Comparison to published data

##### -TOP2i DEG overlap

We obtained a set of DEGs associated with 24 hours of alternative TOP2i treatments (Daunorubicin (DNR), Epirubicin (EPI), Mitoxantrone (MTX)) from Matthews *et al*. (Matthews *et al*. 2024). Of these, 7,703 genes were expressed in our data. In addition, we obtained 278 DEGs associated with CX-5641 treatment for 24 hours from Paul *et al*. (Paul 2025). Of these, 273 were expressed in our data. The proportion of tx+0 DEGs in each set of TOP2i DEGs was determined. Permutation testing (10,000 iterations) was performed to identify whether the overlap of tx+0 DEGs was greater than expected by chance using a random sample of expressed genes in our dataset. Significance was determined following multiple testing correction using the Benjamini-Hochberg method. An adjusted *P* < 0.05 is considered significant.

##### -DNA damage response gene expression analysis

We obtained a set of 66 DNA damage-associated genes from the Molecular Signatures Database (MSigDB v7.5.1) (Liberzon *et al*. 2015). 65 genes are expressed in our data. The proportion of DNA damage response genes in each response cluster was determined. A Fisher’s exact test was used to compare proportions across correlation motif clusters. *P* < 0.05 is considered significant.

##### -p53 target gene expression analysis

We obtained a set of 346 p53 target genes (Fischer 2017). 300 genes are expressed in our data. The proportion of p53 target genes in each response cluster was determined. A Fisher’s exact test was used to compare proportions across response clusters. *P* < 0.05 is considered significant.

##### -DIC gene expression analysis

We compiled a set of 201 unique genes associated with DIC through CRISPR/Cas9 screening (Sapp *et al*. 2021; Liu *et al*. 2024) and *in vitro* functional validation of DIC GWAS genes (Fonoudi *et al*. 2024). Of these, 152 were expressed in our data. The number of overlapping genes in each response cluster was determined.

##### -Heart failure risk gene analysis

Trait associations were downloaded from the NHGRI-EBI GWAS Catalog (Cerezo *et al*. 2025) on 11/13/2025 for heart failure (EFO_0003144), resulting in a list of 299 unique HF GWAS genes. We intersected these genes with our full list of 14,319 expressed genes, determining that 236 genes were expressed in our data, The number of overlapping HF GWAS genes in each response cluster was determined.

All custom scripts are available at https://github.com/mward-lab/Pfortmiller_DOX_recovery_2026 made possible by the workflowr package (Blischak *et al*. 2019).

## Results

### DOX induces transient DNA damage in cardiomyocytes

We differentiated iPSCs from three male and three female healthy donors into iPSC-CMs (Figure 1A). We metabolically selected iPSC-CMs using lactate-containing media and matured the cells in glucose-free CMM media from day 20 to day 30 to promote oxidative phosphorylation, the primary metabolic pathway in adult cardiomyocytes (Rana *et al*. 2012). Flow cytometry analysis of cardiac troponin T (TNNT2) expression indicated a high proportion of cardiomyocytes across individuals (median purity of 91.8%; Figure 1B). In order to investigate the direct effects of DOX treatment on iPSC-CMs over time, we first identified a sub-lethal drug concentration. To do so, we treated iPSC-CMs from three individuals with a range of DOX concentrations (0.01 μM to 1,000 μM) and a DMSO vehicle (VEH) control for 24 hours and determined the proportion of viable cells. We observed a dose-dependent decrease in viability following DOX treatment as expected (median LD50 = 10.09 μM; Figure 1C). We selected a sub-lethal concentration of 0.5 μM DOX (greater than 90% viability following treatment across three individuals) for all subsequent experiments.

**Figure 1:**
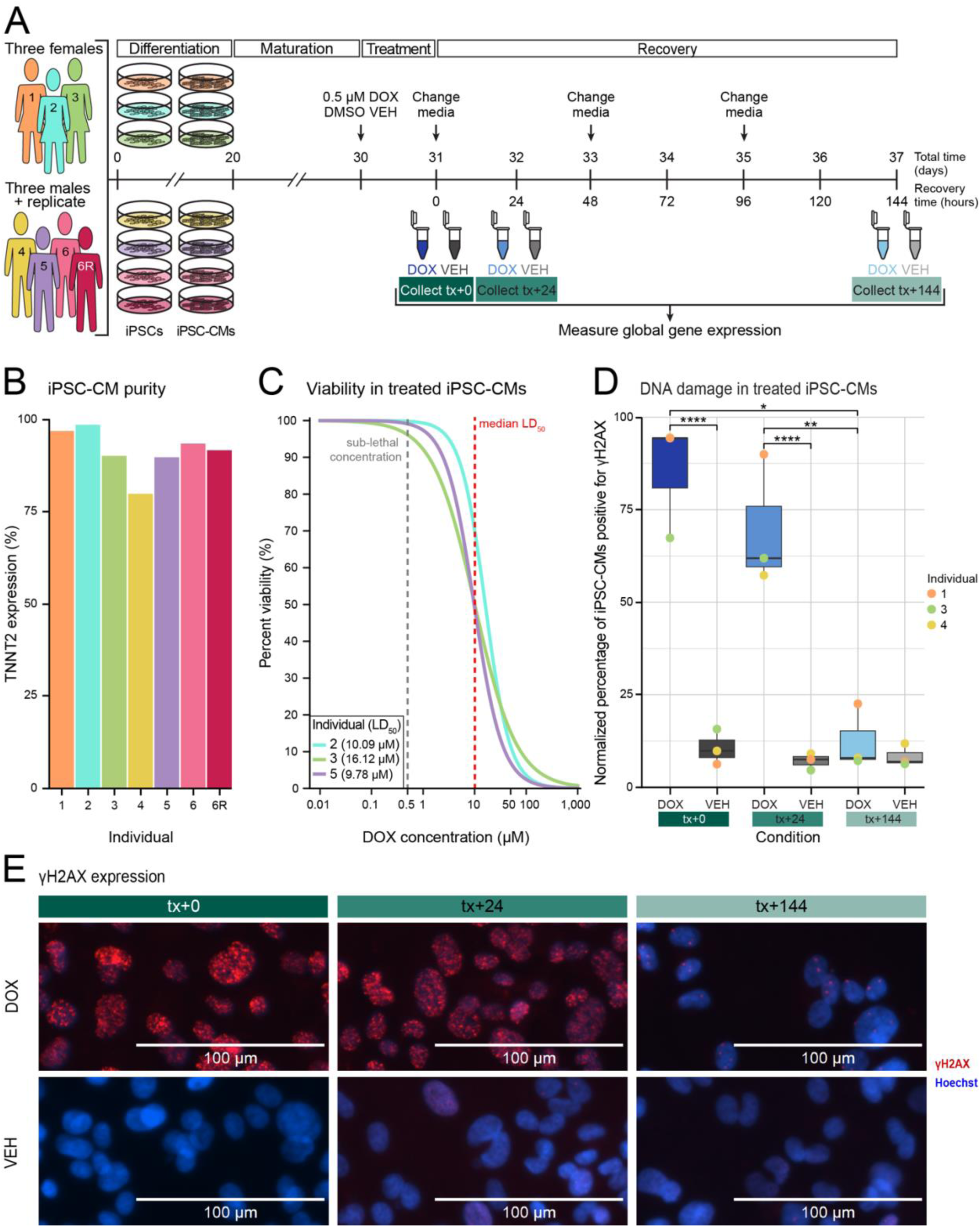
DOX induces DNA damage in iPSC-derived cardiomyocytes. **(A)** Experimental design of this study. iPSCs derived from three females and three males aged between 18-32 were differentiated into cardiomyocytes (iPSC-CMs). iPSC-CMs were treated for 24 hours with 0.5 μM Doxorubicin (DOX) and a Vehicle control (VEH) then assayed directly following treatment (tx+0), after 24 hours of recovery (tx+24), and after 144 hours of recovery (tx+144). **(B)** Purity of differentiated iPSC-CMs as measured by flow cytometry for cardiac troponin T (TNNT2) expression. **(C)** Percent viability of iPSC-CMs over increasing concentrations of DOX after 24 hours of treatment. Cell viability after DOX treatment was assessed in three individuals (Individuals 2,3,5) and normalized to matched VEH. Dose-response curves were generated using a four-point log-logistic regression with the upper asymptote set to 1. LD_50_ values were calculated for each individual, with the median LD_50_ shown in red. The selected sub-lethal concentration for further experimentation is represented by the grey line. **(D)** Quantification of DNA damage as measured by γH2AX expression intensity in DOX- and VEH-treated iPSC-CMs from three individuals (Individuals 1, 3, 4). Asterisks represent a statistically significant difference in γH2AX expression between DOX- and VEH-treated cells at each timepoint, as well as a statistically significant difference between DOX-treated cells across timepoints (\**P* < 0.05, \*\**P <* 0.01, \*\*\**P* < 0.001, t-test). **(E)** Immunofluorescence staining for the γH2AX marker (red) in iPSC-CM nuclei stained with Hoechst (blue) at tx+0, tx+24, and tx+144 in DOX- and VEH-treated cells in a representative individual (Individual 2). Scale bar of 100 μm.

To measure the effect of DOX on cardiomyocytes over time, we treated iPSC-CMs with 0.5 μM DOX and VEH for 24 hours and assayed responses at three timepoints following treatment: i) immediately prior to the removal of DOX and VEH treatment (tx+0), ii) 24 hours post treatment removal (tx+24), and iii) 144 hours post treatment removal (tx+144). Given the known role of DOX in inducing DNA damage, we first asked whether DNA damage is induced in this system, and second whether it is sustained following the removal of DOX. We quantified the effects of DOX on DNA damage by assaying the expression of the DNA double-stranded break marker γH2AX by immunofluorescence staining in three individuals. As expected, at tx+0 there is an increase in DNA damage in DOX-treated iPSC-CMs compared to VEH (t-test; *P* < 0.001; Figure 1D-E). The DNA damage is maintained at tx+24 (*P* < 0.001). However, by tx+144 there is no difference in the level of DNA damage between the DOX- and VEH-treated cells. These results suggest that our model captures DOX-induced DNA damage that clears over time.

### Transcriptional response to DOX decreases over time

We have previously shown that 0.5 μM DOX induces thousands of gene expression changes following 24 hours of treatment in iPSC-CMs (Matthews *et al*. 2024). We therefore asked whether these changes are sustained over time. To do so, we treated iPSC-CMs from six individuals with 0.5 μM DOX and VEH and measured global gene expression by RNA-seq at tx+0, tx+24 and tx+144. We replicated the differentiation and treatment for Individual 6 to account for any unknown technical variation. We processed the 42 samples in treatment-balanced batches across individuals (Table S1). All samples contained at least 20 million reads (Figure S1A), with mapping rates > 99.0% (Table S2). The read depth is similar across treatments (Figure S1B) and timepoints (Figure S1C). We removed lowly expressed genes resulting in a total of 14,319 expressed genes for further analysis. We accounted for unknown technical variation in the data using RUVs and the replicated Individual 6 (Risso *et al*. 2014). Following removal of one factor of unknown variation, principal component analysis reveals that DOX treatment time is associated with the primary source of variation in the data (40.62%; Figure 2A and Figure S2). Similarly, samples cluster by treatment group, and those within the DOX treatment group are clustered by treatment time, unlike the VEH group (Figure S3).

**Figure 2:**
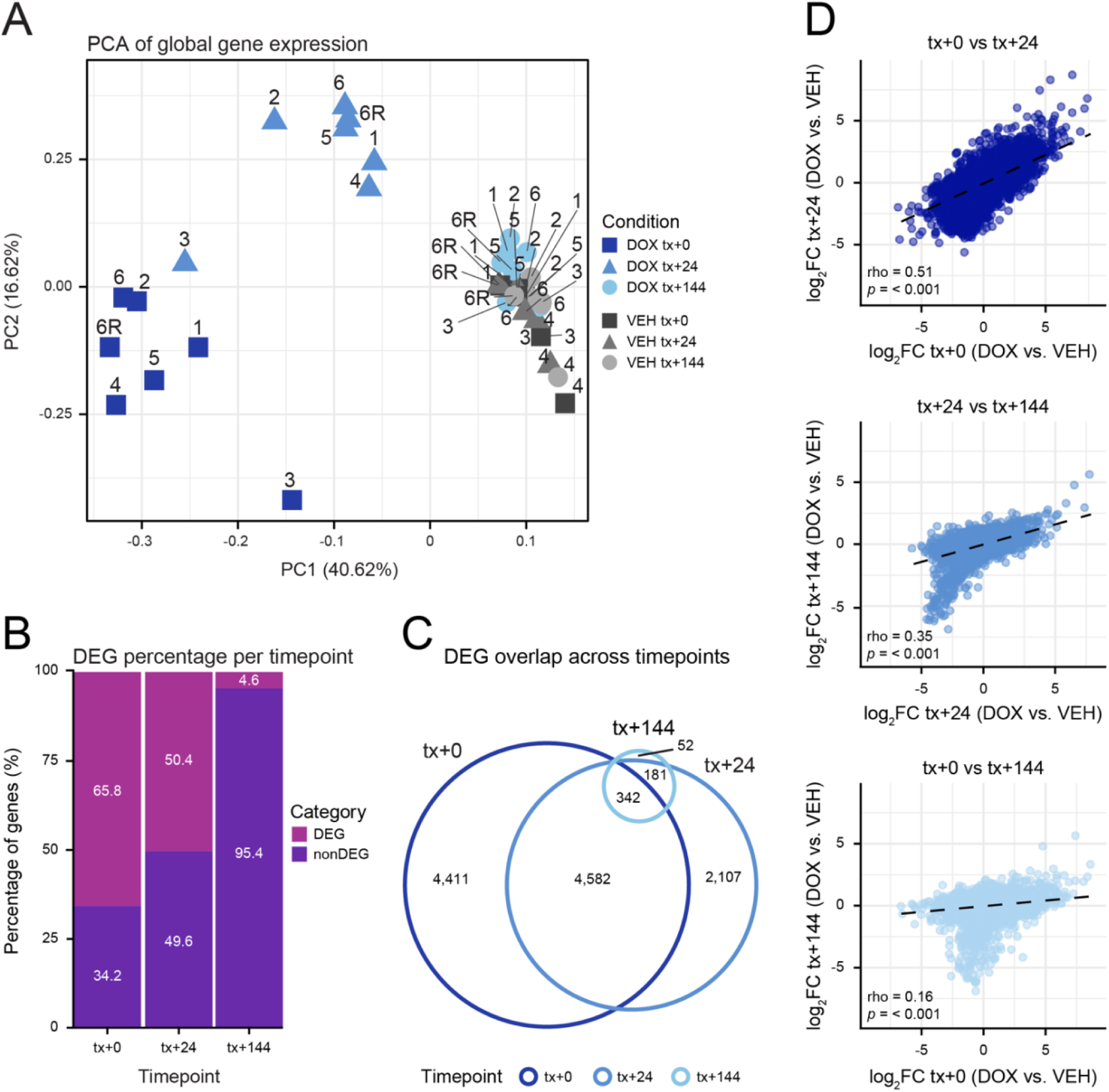
DOX induces global gene expression changes over time. **(A)** Principal component analysis of RUVs-corrected gene expression measurements across 42 samples from six individuals (1, 2, 3, 4, 5, 6) with a replicate of individual 6 (6R) at treatment timepoints (DOX_tx+0: dark blue, DOX_tx+24: medium blue, DOX_tx+144: light blue, DMSO_tx+0: dark grey, DMSO_tx+24: medium grey, DMSO_tx+144: light grey). Data is representative of log_2_ cpm values of 14,319 expressed genes. **(B)** Proportion of differentially expressed genes (DEGs) between DOX- and VEH-treated samples out of 14,319 expressed genes across timepoints. **(C)** Overlap of DEGs at each timepoint. **(D)** Spearman correlation (ρ) of log_2_ fold change response magnitude between timepoints.

To identify DOX response genes at each timepoint, we performed pairwise differential expression analysis by comparing DOX and VEH treatments. We randomly selected one Individual 6 replicate and included the RUVs-identified weight as a covariate in the linear model. We find thousands of differentially expressed genes (DEGs) at tx+0 and tx+24, and hundreds at tx+144 (DOX vs VEH tx+0 = 9,420; tx+24 = 7,212; tx+144 = 660; 5% FDR; Figure S4; Table S3-5). These changes correspond to 66% of all expressed genes being DEGs in tx+0, 50% in tx+24 and 5% in tx+144 (Figure 2B). While there is a roughly even distribution of DEGs that are up- and down-regulated at tx+0 and tx+24 (48% upregulated for both), the majority of tx+144 DEGs are downregulated (71%; n = 466; Figure S4).

We then asked how many of the DEGs are shared across timepoints. We find that nearly half of all tx+0 DEGs (49%; n = 4,582) are DEGs in tx+24, and only 342 (4%) are DEGs in tx+0, tx+24 and tx+144 (Figure 2C). To compare global responses across conditions, we considered the magnitude of the treatment effect (log_2_ fold change) across all genes rather than just those that meet an FDR-based cutoff. There is a strong correlation between responses at early timepoints (tx+0 vs tx+24 ρ = 0.51; *P* < 0.001), which decreases over time (tx+24 vs. tx+144 ρ = 0.35; *P* < 0.001; tx+0 vs tx+144 ρ = 0.16; *P* < 0.001 Figure 2D). These findings suggest that most genes return to baseline expression levels seven days after DOX treatment.

### Hundreds of response genes are sustained days after DOX treatment

To determine gene expression response dynamics over time, we used a Bayesian approach to jointly model pairs of tests at each timepoint (See Methods). Following Bayesian information criterion (BIC) and Akaike information criterion (AIC) analysis, we find that the major expression patterns are explained by two response clusters (Figure S5). The first cluster corresponds to genes that respond only in tx+0 and not in tx+24 or tx+144, which we define as ‘acute response’ genes (Figure 3A; Table S6). The second cluster corresponds to genes that respond in all three timepoints, which we define as ‘chronic response’ genes. The majority of genes assigned to a cluster are classified as acute response genes (n = 7,602; 94% of 8,103 assigned genes). However, there are 501 chronic response genes. The absolute log_2_ fold change of genes in the acute response cluster approaches zero over time, while genes in the chronic response cluster do not (Figure 3B). In fact, many of the chronic response genes increase in effect over time (Figure S6). Individual genes assigned to these clusters show the expected expression patterns. For example, *ZMYND8* is a gene in the acute response cluster that is downregulated specifically at tx+24 (Figure 3C). *CCNB2* in the chronic response cluster is increasingly downregulated across timepoints (Figure 3D).

**Figure 3:**
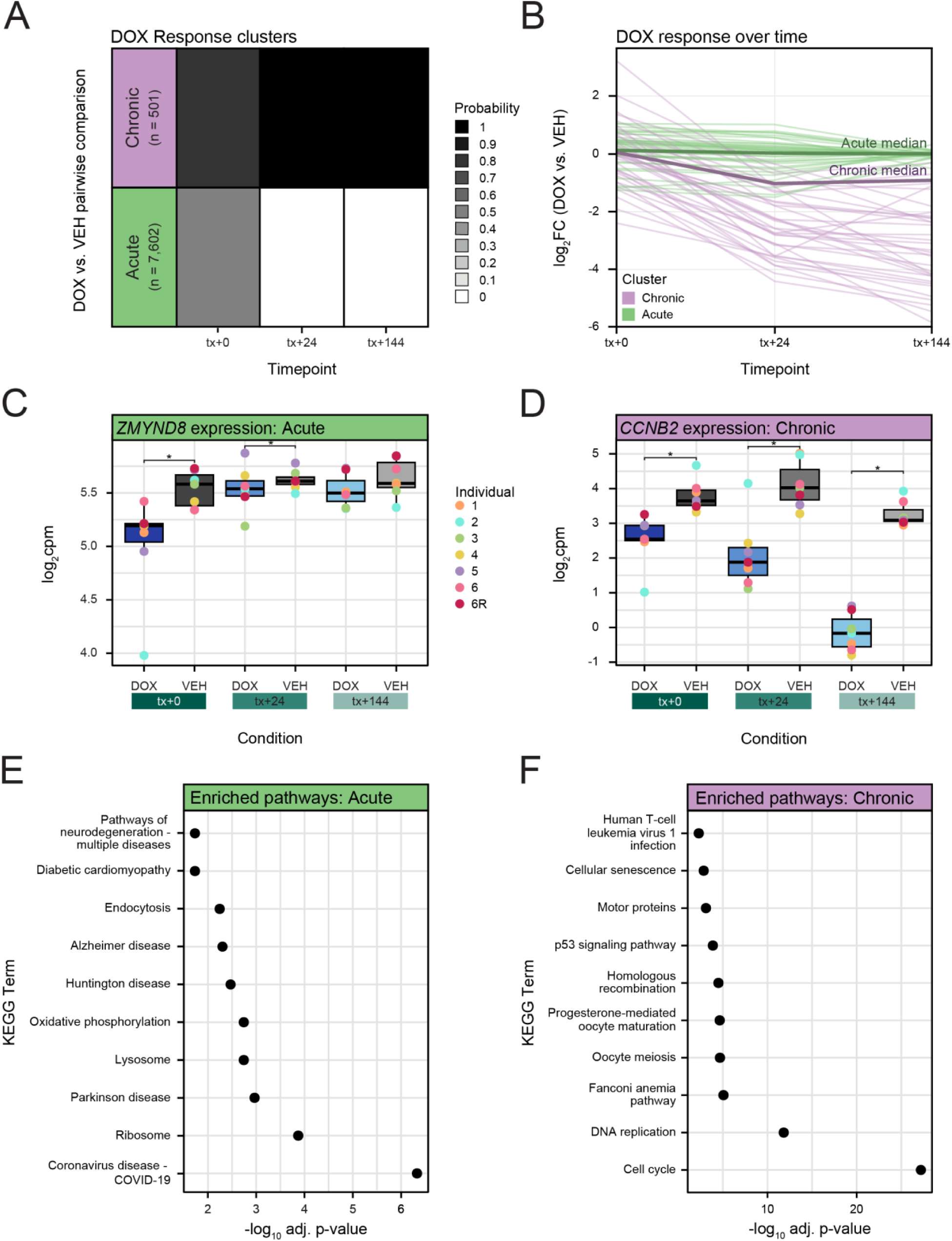
Trajectory analysis reveals acute and chronic DOX response genes. **(A)** Gene expression clusters identified through joint modeling of test pairs over time. Shades of grey represent the posterior probabilities of genes being differentially expressed at each timepoint. Genes are categorized into two clusters based on their likelihood of belonging to a cluster and posterior probabilities: genes with probability >= 0.3 – 0.6 in tx+0 are designated as ‘acute response’ (green), while those with probability > 0.5 in every timepoint are designated as ‘chronic response’ (purple). **(B)** Absolute log_2_ fold change of the top 100 genes from acute (green) and chronic (purple) response clusters. Median log_2_ fold change (bold line) for each cluster is shown. **(C)** Gene expression (log_2_ cpm) of the *ZMYND8* acute response gene over time. Asterisks represent timepoints where the gene is classified as a DEG. **(D)** Gene expression (log_2_ cpm) of the chronic response gene *CCNB2* over time. **(E)** Biological pathways enriched in acute response genes. The top 10 KEGG terms that are significantly enriched in the acute gene set compared to all expressed genes are shown. **(F)** Top 10 KEGG terms significantly enriched amongst chronic response genes.

To identify biological pathways associated with each response cluster, we performed pathway enrichment analysis. The acute response cluster is enriched for terms related to the ribosome, and lysosome as well as oxidative phosphorylation and endocytosis compared to all expressed genes (Fisher’s one-tailed test; adj. *P* < 0.05; Figure 3E). The chronic response cluster is enriched for cell cycle (*CDKN1A*, *CDC20*, *E2F2)*, DNA replication (*POLD3*, *FEN1*, *MCM2*), p53 signaling (*CCNB2*, *FAS*, *CDK1*), homologous recombination (*RAD51*, *BRCA1*, *BLM*), and cellular senescence pathways (*CDK1*, *CHEK1*, *MAP2K6*; adj. *P* < 0.05; Figure 3F). This suggests that the two response clusters represent distinct sets of genes associated with distinct cellular processes.

### Chronic response genes respond to treatment with other TOP2i

As the primary mechanism of action of DOX is associated with inhibition of TOP2, we asked whether the response patterns we observe are recapitulated across other TOP2i drugs. We obtained DEGs from iPSC-CMs treated for 24 hours with 0.5 μM TOP2i. 51.9 – 83.9% of tx+0 DEGs are DEGs following DOX treatment in other studies (Matthews *et al*. 2024; Paul 2025). The higher overlap set was from a study which included the same panel of individuals, while the lower overlap set had only three individuals in common and included female individuals only. As expected, anthracyclines showed the highest overlap with tx+0 DEGs (54.5% for Daunorubicin and 52.5% for Epirubicin), followed by the anthracenedione Mitoxantrone (8.0%; Figure 4A). The unrelated G-quadruplex inhibitor CX-5461 exhibited the lowest overlap (1.7%).

**Figure 4:**
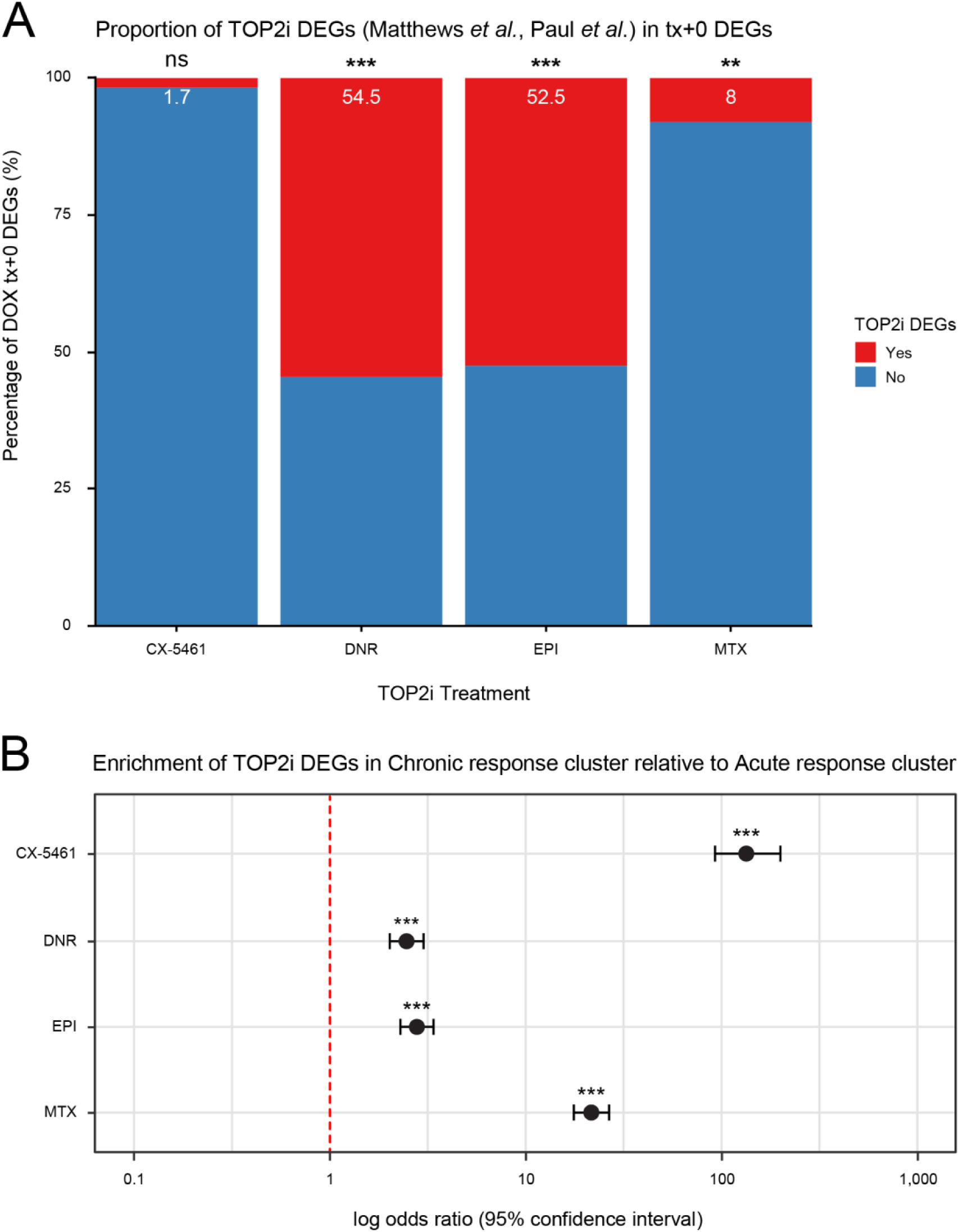
Chronic response genes are enriched in TOP2i DEGs compared to acute response genes. **(A)** Proportion of tx+0 DEGs that overlap DEGs following 24 hours of treatment with other TOP2i (red). Data was obtained for CX-5461 from Paul *et al*. (Paul 2025) and Daunorubicin (DNR), Epirubicin (EPI) and Mitoxantrone (MTX) from Matthews *et al*. (Matthews *et al*. 2024). Significance of the overlap was determined by permutation testing (\*\*\**P* < 0.001). **(B)** Enrichment of TOP2i DEGs in chronic response genes compared to acute response genes. A Fisher’s exact test was used to calculate enrichment where Odds ratios and 95% confidence intervals are plotted. Asterisk represents a significant enrichment (\*\*\**P* < 0.001).

We then asked whether the chronic response genes are likely to be relevant across TOP2i. Indeed, we find that the majority of chronic response genes respond to other TOP2i (68.7% respond to Daunorubicin, 65.9% to Epirubicin, 56.9% to Mitoxantrone and 40.5% to CX-5461; Figure S7). Notably there is an enrichment of all TOP2i DEGs in the chronic response cluster compared to the acute response cluster (Fisher’s exact test; *P* < 0.05; Figure 4B). These results suggest that chronic response genes, that do not recover from DOX treatment, may be relevant to TOP2i more broadly.

### Chronic response genes are enriched for DNA damage-associated genes

Given that we observed DOX-induced DNA damage that was restored over time at the cellular level, we asked whether genes known to be relevant to the response to DNA damage are amongst the chronic response gene set. First, we obtained a set of 66 genes associated with the DNA damage response (DDR) (Liberzon *et al*. 2015) and calculated the proportion of DDR genes in the acute and chronic response clusters. We found that the chronic response cluster is enriched for DDR genes compared to the acute response cluster (Fisher’s exact test; OR = 10.4, confidence interval (CI) = 5.1 – 20.5; *P* < 0.001; Figure 5A). Second, we obtained a set of 346 genes identified as transcriptional targets of the DNA damage-associated transcription factor p53 (Fischer 2017). Again, we found that the chronic response cluster is enriched for p53 target genes compared to the acute response cluster (Fisher’s exact test; OR = 2.5, confidence interval (CI) = 1.6 – 3.9; *P* < 0.001; Figure 5A).

**Figure 5:**
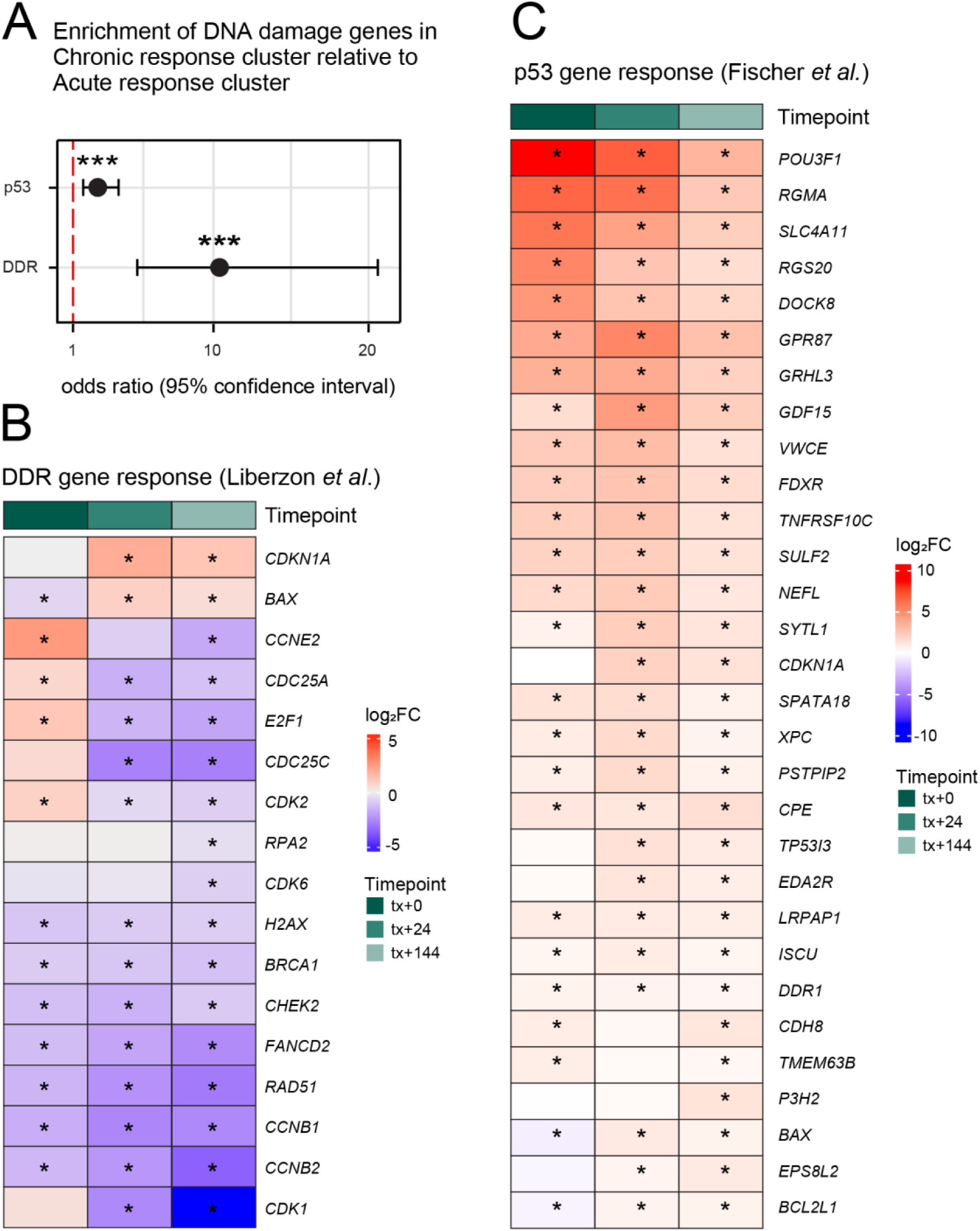
Chronic response genes are enriched for DNA damage-associated genes. **(A)** Enrichment of 66 DDR genes (Liberzon *et al*. 2015) and 346 p53 target genes (Fischer 2017) in chronic response genes compared to acute response genes. Asterisk represents a significant enrichment as calculated by Fisher’s exact test (\*\*\**P <* 0.001). Odds ratios and 95% confidence intervals are plotted. **(B)** Gene expression response (log_2_ fold change) across timepoints of DDR genes that are classified as DEGs at tx+144. Color scale represents either upregulation (red), no change (white), or downregulation (blue). Asterisks represent genes that are classified as DEGs in each respective timepoint. **(C)** Gene expression response (log_2_ fold change) across timepoints of p53 response genes that are DEGs at tx+144.

In line with DNA damage-associated genes being enriched in the chronic response cluster, 11 of 65 DDR genes are classified as DEGs in all timepoints including *CCNB2*, *FANCD2* and *RAD51* that relate to cell cycle and repair pathways (Figure 5B). DDR genes are enriched in tx+144 DEGs compared to non-DEGs at this timepoint (Fisher’s exact test; OR = 7.5; 95% confidence interval (CI) = 4.0-13.3; *P* < 0.05; Figure S8A); however DDR genes are not enriched in the earlier timepoints. We also find that many p53 target genes are DEGs in each timepoint (82 of 300) including *NEFL*, *XPC* and *BAX* (Figure 5C). p53 target genes are enriched in DEGs at all timepoints (Fisher’s exact test; tx+0: OR = 1.5; 95% confidence interval (CI) = 1.1-1.9; tx+24: OR = 2.1, CI = 1.6-2.8; tx+144: OR = 2.3; CI = 1.5-3.4; *P* < 0.05; Figure S8B). Together, these results suggest that the genes that do not recover from DOX treatment relate to DNA damage in line with them being enriched in p53 and cell cycle associated pathways.

### Over one hundred genes show increased response to DOX over time

As we noted above, many chronic response genes show increased response to DOX over time. We therefore further stratified the 501 genes within the chronic response cluster to isolate the set where the response to DOX increases over time (absolute log_2_ fold change tx+0 < absolute log_2_ fold change tx+24 < absolute log_2_ fold change tx+144; Table S7). This yielded 163 progressively chronic response genes (Figure 6A). 94% of these genes are progressively downregulated. These genes are enriched for cell cycle (e.g. *CDC20*, *ORC1*, *KNL1*) and DNA repair processes (*RAD51*, *EXO1*, *MCM7*) (Figure 5B). This is in contrast to genes that are chronic, but do not increase in response over time, that are uniquely enriched for processes related to ligand-receptor interactions (*TNFRSF9*, *ITGA8*, *EDNRB*), p53 (*FAS*, *CHEK1*, *CDKN1A*) and PI3K-Akt signaling (*NGF*, *CDK2*, *FGFR3*) (Figure 5C-D).

**Figure 6:**
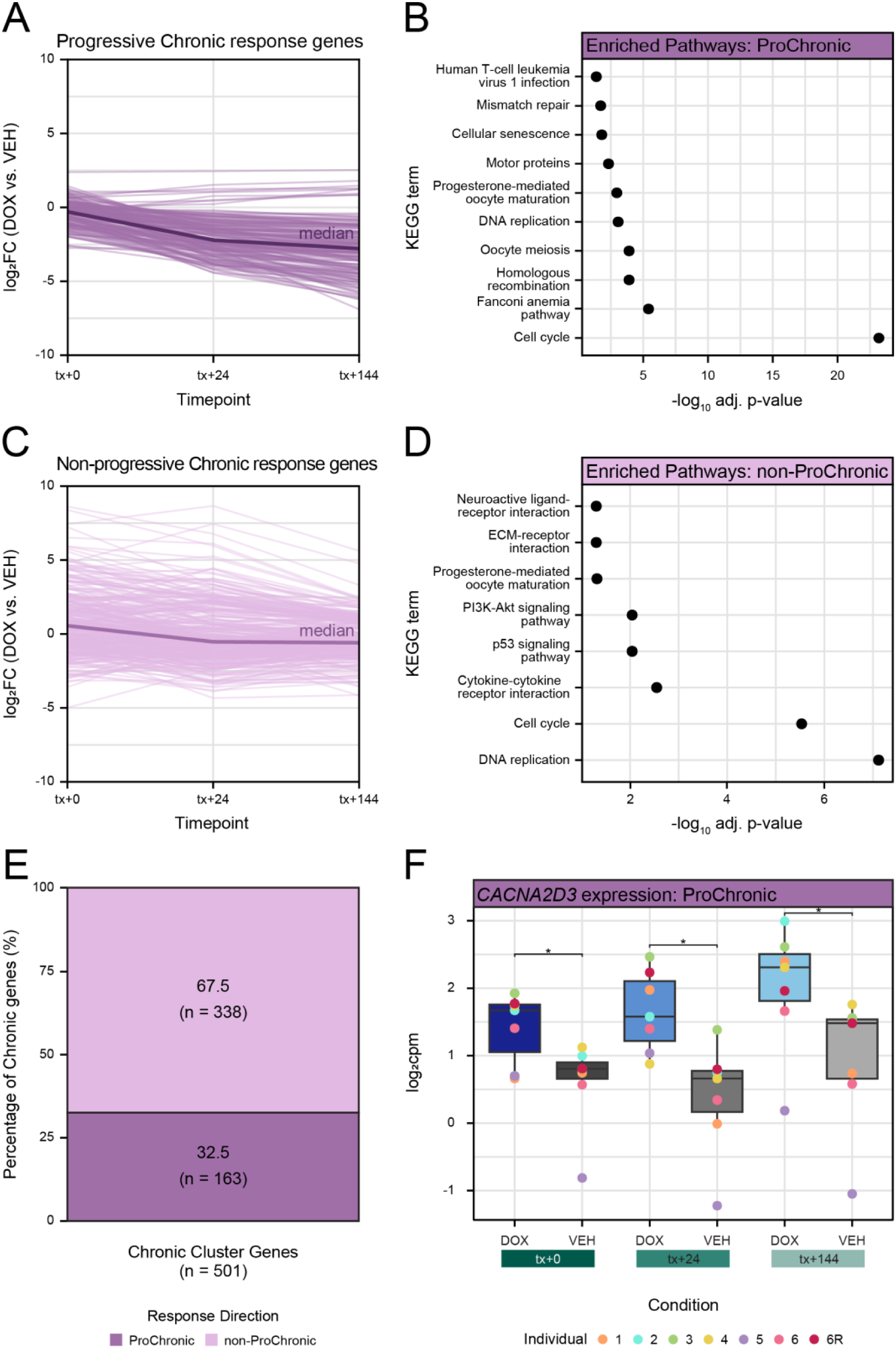
Over a hundred genes continue to diverge from baseline seven days post treatment. **(A)** Absolute log_2_ fold change of chronic response genes that continue to diverge from baseline expression over time (progressively chronic). Median log_2_ fold change (bold line) is shown. **(B)** Biological pathways enriched in progressively chronic response genes. The top 10 KEGG terms that are significantly enriched in the acute gene set compared to all expressed genes are shown. **(C)** Absolute log_2_ fold change of chronic response genes that do not continue to diverge from baseline expression over time. Median log_2_ fold change (bold line) is shown. **(D)** Biological pathways enriched in chronic response genes that do not progressively increase in response. The top 10 KEGG terms that are significantly enriched in this gene set compared to all expressed genes are shown. **(E)** Proportion of chronic response genes that are progressively chronic and not. **(F)** Gene expression (log_2_ cpm) of the *CACNA2D3* progressively chronic response gene over time. Asterisks represent timepoints where the gene is classified as a DEG.

There are nine genes that are progressively upregulated. These include *ASTN1*, *EPS8L2*, *CEMIP*, *P3H2*, *CACNA2D3*, *PDE11A*, *ACHE*, *CPE* and *PTCHD4*. *CACNA2D3* is the target of amlodipine, an approved drug for hypertension and angina. Blocking *CACNA2D3* with amlodipine has been shown to inhibit DOX-induced apoptosis in neonatal rat cardiomyocytes (Yamanaka *et al*. 2003). These results suggest that there are a small number of genes that continue to diverge in expression from baseline seven days following treatment.

### Most DIC genes are acute response genes

DOX treatment is associated with DIC in some patients. GWAS has identified tens of genetic variants associated with risk for developing cardiotoxicity following DOX treatment (Schneider *et al*. 2017; Park *et al*. 2020). 25 genes in these loci have been functionally-validated in *in vitro* models (Fonoudi *et al*. 2024). Large-scale CRISPR/Cas9 screens in iPSC-CMs have also identified genes which mediate the effects of DOX (Sapp *et al*. 2021; Liu *et al*. 2024). We therefore combined the sets of genes identified by these studies into a DIC gene set consisting of 201 genes expressed in our iPSC-CMs, and investigated their longitudinal response to DOX. Of the 81 genes that are classified in response signatures, 94% are classified as acute response genes (n = 76; Figure 7A). One gene is classified as progressively chronic, *POLQ*. This gene, which encodes polymerase theta, is important in rapid, but low fidelity DNA repair (Wood and Doublie 2016). Its expression is progressively downregulated over time (Figure 7B). Four genes are classified in the broader chronic response set: *CPMK2*, *HUNK*, *TAGLN2* and *COL19A1*.

**Figure 7:**
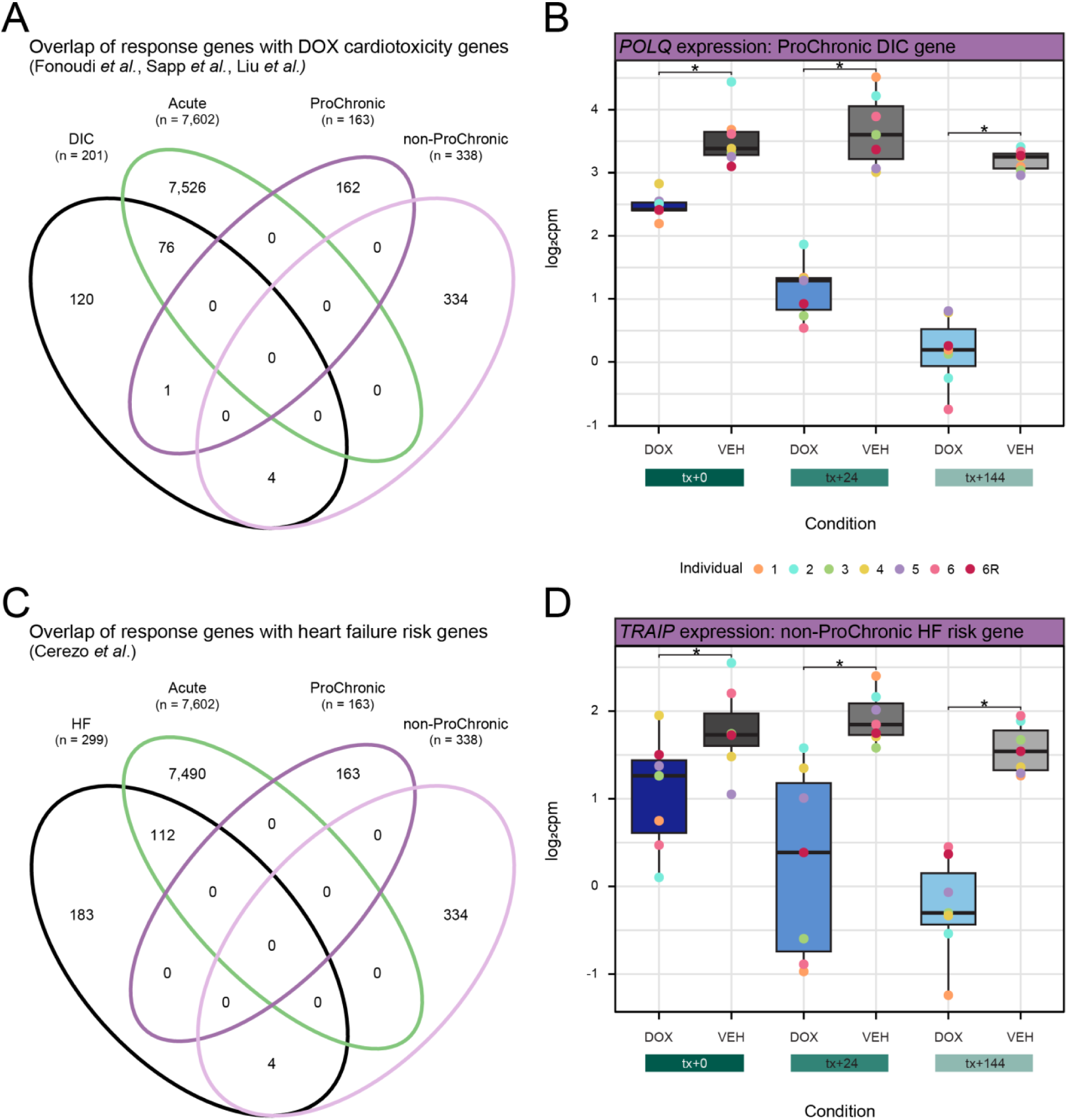
Chronic DOX response genes are relevant to DIC and heart disease. **(A)** Overlap between DIC cardiotoxicity genes (functionally-validated DOX targets (Fonoudi *et al*. 2024), and CRISPRi targets (Sapp *et al*. 2021; Liu *et al*. 2024)) and acute, chronic and progressively chronic response genes. **(B)** Gene expression (log_2_ cpm) of the DIC gene *POLQ* that is classified as a progressively chronic response gene over time. Asterisks represent timepoints where the gene is classified as a DEG. **(C)** Overlap between heart failure risk loci genes (Cerezo *et al*. 2025) and acute, chronic and progressively chronic response genes. **(D)** Gene expression (log_2_ cpm) of the heart failure gene *TRAIP* that is classified as a chronic response gene over time.

Given that DIC can lead to heart failure, we obtained the set of genes in HF risk loci (Cerezo *et al*. 2025). Of the 116 genes that are classified as response genes, 97% are classified as acute response genes (n = 112; Figure 7C). There are no progressively chronic genes associated with heart failure but four including *ABCA12*, *CDKN1A*, *CIT* and *TRAIP*, are in the chronic set that is not progressively different over time. *TRAIP* is downregulated in response to DOX and is known to be involved in the DNA damage response (Figure 7D) (Harley *et al*. 2016).

## Discussion

DIC is a well-established phenomenon in cancer patients treated with DOX. Many studies have indicated widespread effects on global gene expression in human cardiomyocytes following acute exposure to DOX and linked these effects to the phenotype in humans (Burridge *et al*. 2016; Knowles *et al*. 2018; Matthews *et al*. 2024). Here, we measured the effects of DOX treatment and recovery from treatment to mimic chronic effects of the drug. We identified thousands of gene expression changes that diminish over time. However, we also identified a chronic response signature corresponding to p53 activation and suppression of DNA repair.

By measuring gene expression trajectories following DOX treatment and removal, we identified two predominant response signatures – genes that respond to treatment and return to baseline levels of expression following 24 hours of recovery, and genes that respond to treatment and sustain their response following six days of recovery. The vast majority of response genes return to baseline during recovery (94%); however there are 501 genes that show perturbed expression throughout the recovery period.

Chronic response genes are enriched for DNA damage response pathways and p53 signaling. P53 target genes continue to be activated over time, while DNA damage response genes are progressively downregulated. However, we observe that DNA damage, as measured by the expression of γH2AX, is diminished at the final timepoint suggesting that double-strand breaks have been resolved. The discrepancy between observed DNA damage and DNA damage response gene mRNA changes may suggest a chronic stress state, or secondary effects of the DOX treatment. For example, there may be oxidative DNA damage initiated through drug-induced reactive oxygen species (Yoshida *et al*. 2009); however classic markers of oxidative DNA damage including *OGG1*, *PARP1* and *ATM* are DEGs in early timepoints only.

Low doses of DOX treatment in cardiomyocytes have been shown to induce a senescent-type state with characteristic beta galactosidase activity (Maejima *et al*. 2008). When considering a set of 55 senescence-associated genes identified from a range of cell types, excluding cardiomyocytes, we find one gene, *PLK3*, to be within our set of chronic response genes (Hernandez-Segura *et al*. 2017). Senescence-associated phenotype markers including *IL6*, *IL8*, *MMP3* and *CXCL1* are not expressed in our cardiomyocytes; however *GDF15* is, and is upregulated at all timepoints. Senescence has been observed in the hearts of patients treated with DOX, and *GDF15* is increased following a 28-day DOX treatment schedule in engineered heart tissue (Linders *et al*. 2023).

We note that the chronic response genes are largely distinct from genes that have previously been associated with DIC through global CRISPRi screening or CRISPRi targeted to genes in DIC-associated loci (Sapp *et al*. 2021; Fonoudi *et al*. 2024; Liu *et al*. 2024). This suggests the identification of a set of genes for further investigation as potential biomarkers or therapeutic targets. Indeed, we identify *CACN2D3* as a gene that is progressively upregulated over time. This gene is important to cardiac function and drugs that target it are used in the treatment of angina.

The limitation of iPSC-derived cardiomyocytes is that they resemble fetal cardiomyocytes more than adult cardiomyocytes. It is possible that the response to DOX, and the DNA repair mechanisms differ between adult cardiomyocytes and iPSC-derived cardiomyocytes. To mitigate this potential effect, we used 30-day iPSC-CMs cultured in media to shift the metabolic state of the iPSC-CMs to be reflective of adult cardiomyocytes. We note however that in humans, cardiomyocytes proliferate up until age 20 (Mollova *et al*. 2013). Given that there are many pediatric cancers that are treated with DOX, we believe that this system is physiologically and clinically relevant.

In summary, we measured the effects of treatment and recovery from DOX in cardiomyocytes from six individuals. We found that DOX induces DNA damage, which is resolved following six days of recovery from treatment. Similarly, large-scale effects on gene expression are induced following treatment that diminish over time. However, there are a set of 501 genes that are persistently perturbed six days following treatment that are related to p53 signaling and DNA damage. Our results suggest that chronic modeling of DOX treatment can highlight genes that may relate to the long-term effects of DOX treatment.

## Acknowledgements

We thank all members of the Ward Lab for helpful discussions particularly Omar Johnson. We thank Kelly Frazer and the University of California San Diego for providing the iPSC lines through the iPSCORE resource. We thank the Next Generation Sequencing Core Facility at the University of Texas Medical Branch for preparing and sequencing the RNA-seq libraries, and the Flow Cytometry Core Facility for access to flow cytometers. The authors acknowledge the Texas Advanced Computing Center (TACC) at The University of Texas at Austin for providing HPC resources that have contributed to the research results reported within this paper (http://www.tacc.utexas.edu). This work was supported by a CPRIT Scholar award to M.C.W.

## Funding

This work was funded by a Cancer Prevention Research Institute of Texas (CPRIT) Recruitment of First-Time Faculty Award (RR190110) to M.C.W. E.M.P was supported by a Jeane B Kempner Pre-doctoral Fellowship administered through UTMB.

## Author contributions

M.C.W conceived and designed the study. E.M.P, A.R.B, J.A.G performed experiments. E.M.P analyzed the data. E.M.P and M.C.W wrote the manuscript with input from co-authors. M.C.W supervised the work.

## Supplemental material

S1 Appendix: Document containing Supplemental Figures 1-8

S2 Appendix: Document containing Supplemental Tables 1-7

## Notes

### Competing Interest Statement

The authors have declared no competing interest.

