## Supplementary material for "Progressive suppression of DNA repair genes with persistent p53 activation in Doxorubicin-treated cardiomyocytes": S1 Appendix

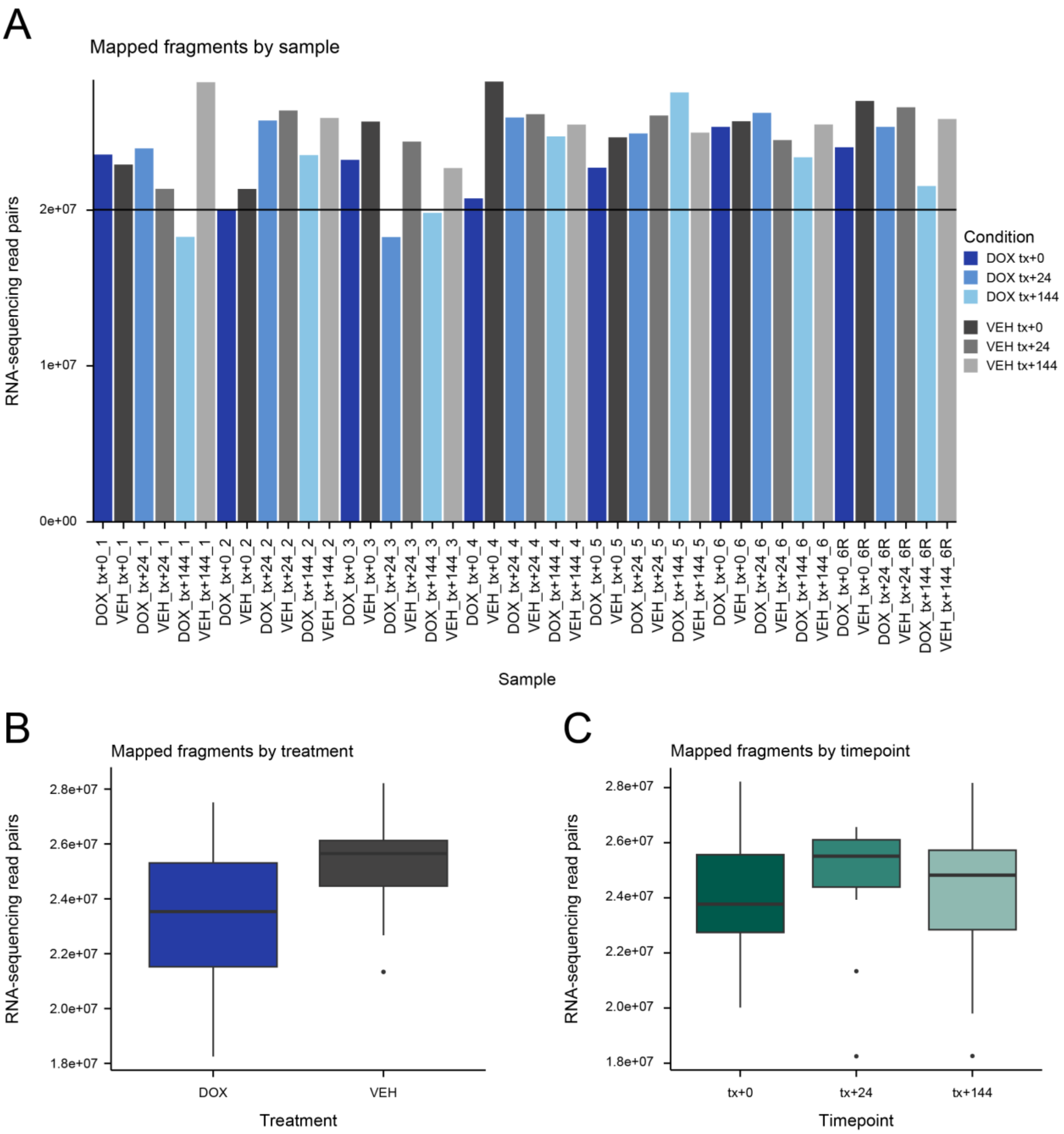


**Supplementary Figure 1: RNA-seq data has a similar number of mapped fragments across treatments and timepoints. (A)** Total number of mapped fragments per sample for 42 DOX- (blue) and VEH-treated (grey) samples across six individuals including a technical replicate of Individual 6. **(B)** Total number of mapped fragments for all samples within a treatment group (DOX: blue, VEH: grey). **(C)** Total number of mapped fragments for all samples within a timepoint group (tx+0: dark green; tx+24: medium green; tx+144: light green).

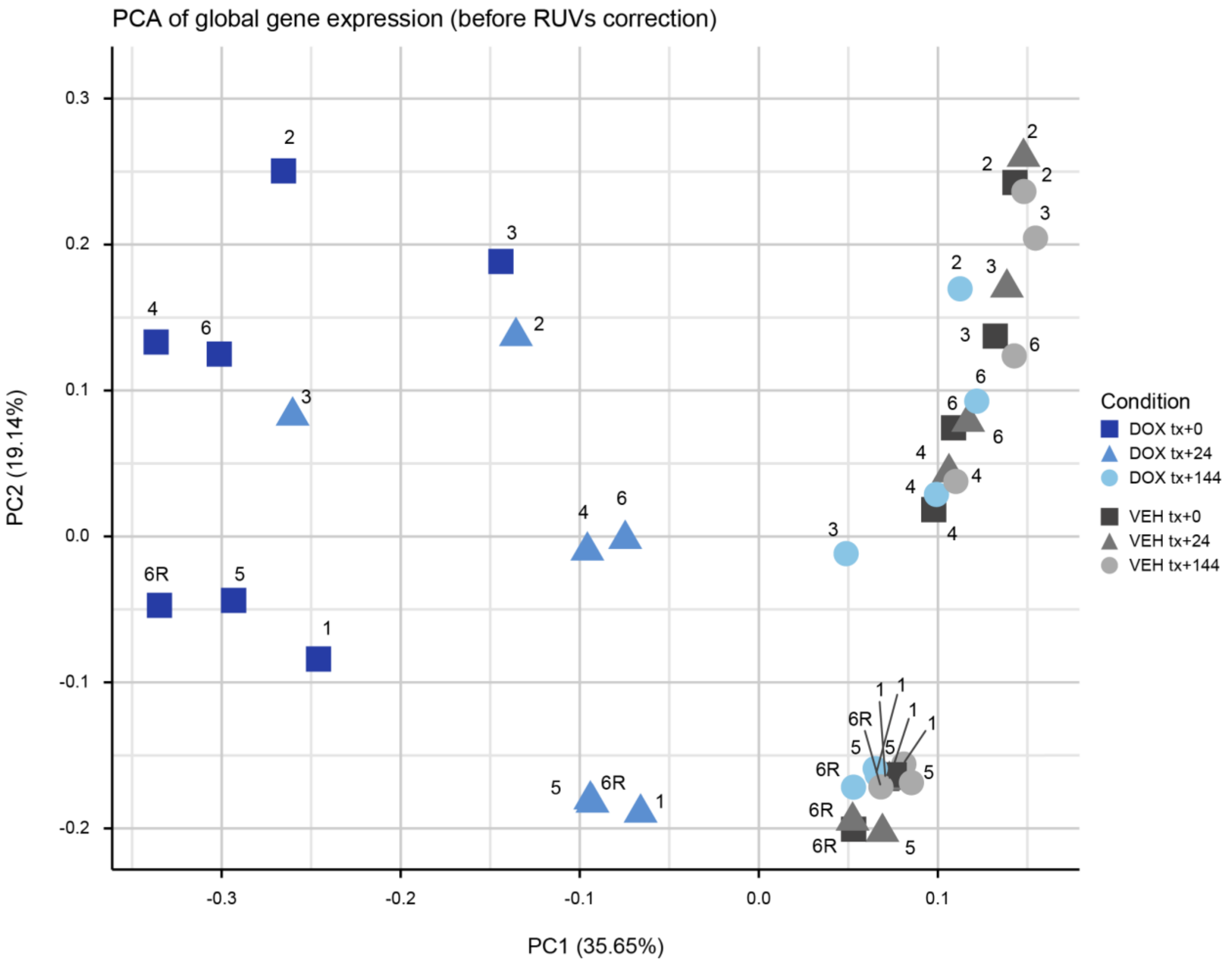


**Supplementary Figure 2: Technical replicates do not cluster together prior to removal of unwanted variation. (A)** Principal component analysis of gene expression ($log_2$ cpm) of 42 RNA-seq samples across 6 individuals (1, 2, 3, 4, 5, 6) with a technical replicate (6R), over three timepoints (tx+0: square; tx+24: triangle; tx+144: circle), in two treatments (DOX: blue; VEH: grey). Values are shown in $log_2$ cpm prior to correction with RUVs.

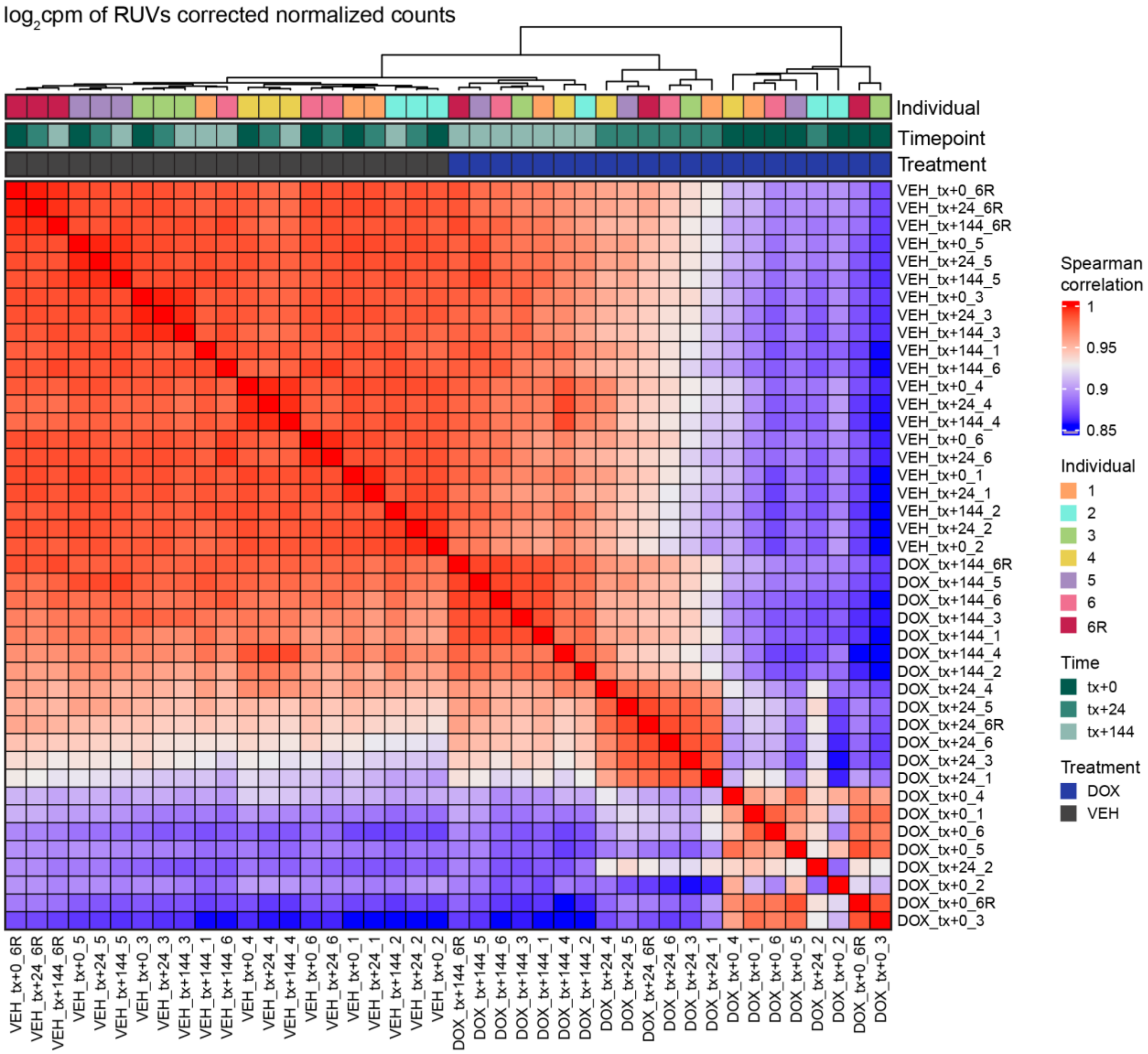


**Supplementary Figure 3: Samples cluster by treatment and then by timepoint.** Correlation heatmap of 42 RNA-seq samples following RUVs correction using a technical replicate. Color scale represents Spearman correlation coefficient. Data represented is the log$_2$ cpm of RUVs normalized counts. Top annotation is of two treatments (DOX: blue; DMSO: grey) across three timepoints (tx+0: dark green; tx+24: medium green; tx+144: light green) in 6 individuals with a technical replicate (1: orange; 2: cyan; 3: green; 4: yellow; 5: purple; 6: pink; 6R: red).

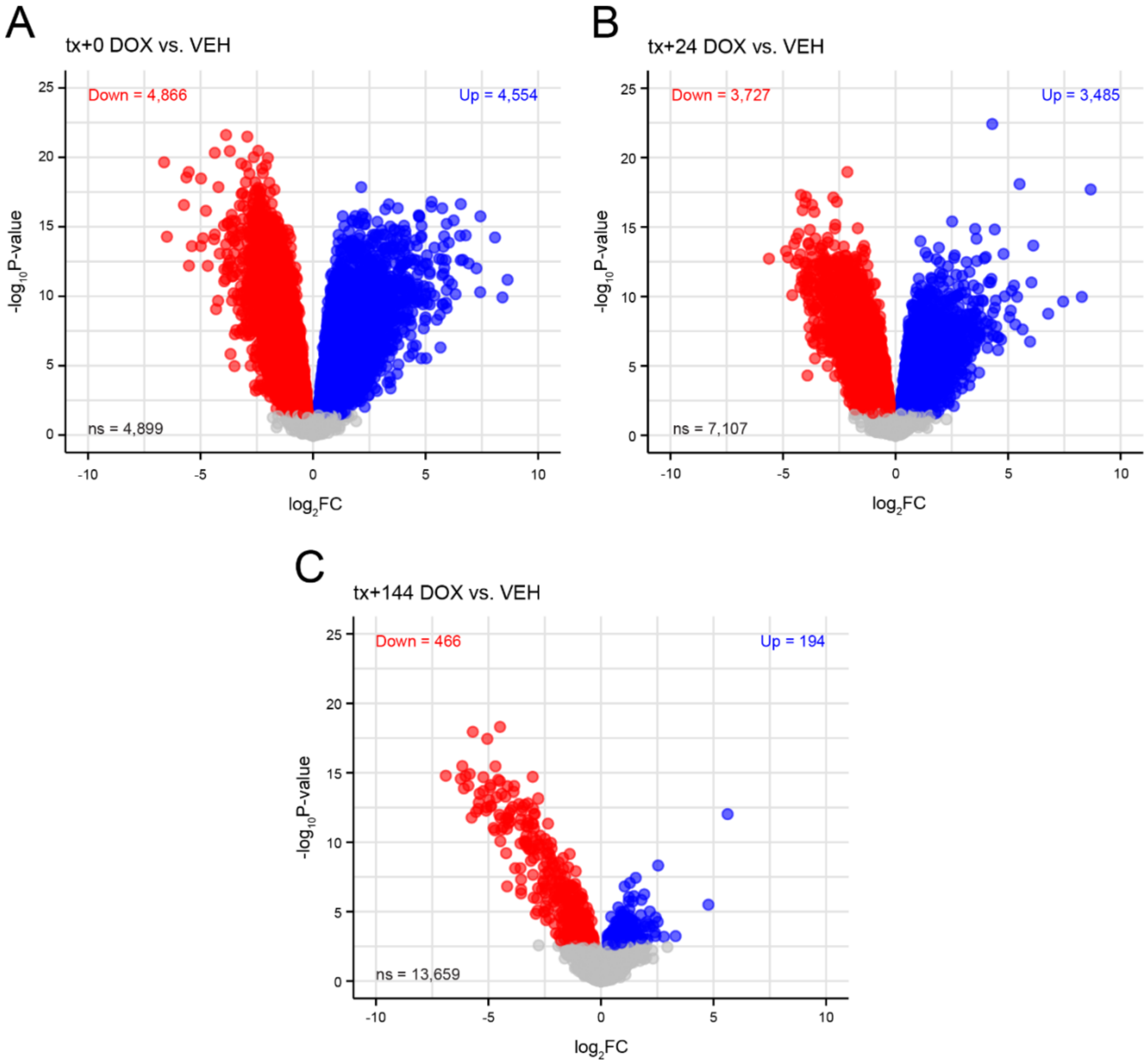


**Supplementary Figure 4: DOX response diminishes over time. (A)** Gene expression response magnitude ($\log_2$ fold change) and significance ($-\log_{10}$ *P*) of all genes expressed at tx+0. Differentially expressed genes (DEGs) are denoted in red if downregulated in response to DOX or blue if upregulated, based on an adj. $P < 0.05$. Genes that are not DEGs are denoted in grey. **(B)** Gene expression response at tx+24 as for (A). **(C)** Gene expression response at tx+144 as for (A).

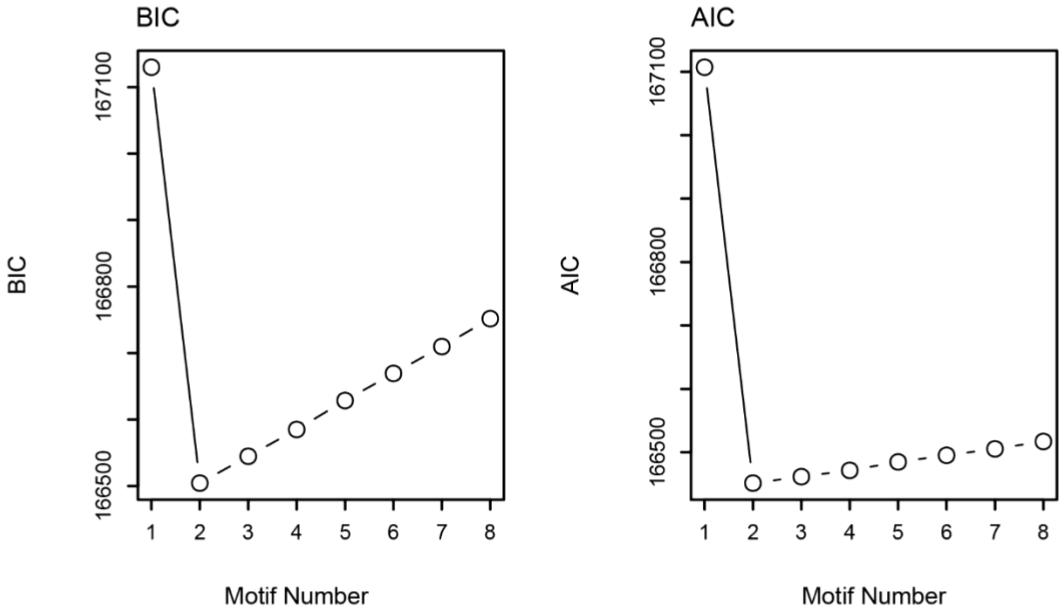


**Supplementary Figure 5: Trajectory analysis identifies two gene response clusters.** Bayesian Information Criterion (BIC) and Akaike Information Criterion (AIC) of jointly modeled test pairs derived from Cormotif, where the lowest point on the graph shows the appropriate number of motifs for these data.

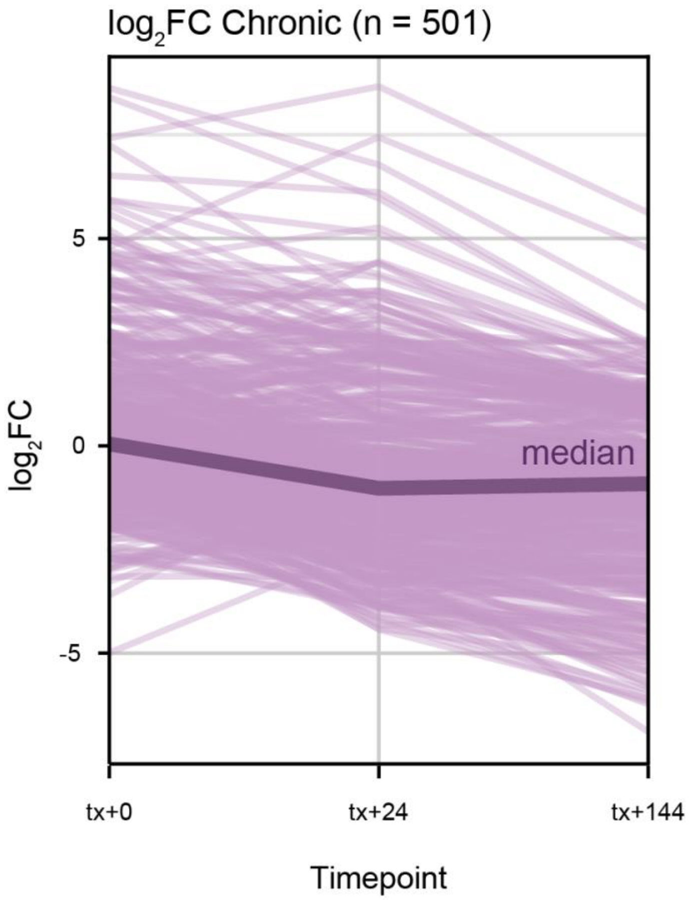


**Supplementary Figure 6: Gene expression response patterns amongst Chronic response genes.** $Log_2$ fold change of all 501 genes from the Chronic response cluster over time (tx+0, tx+24, tx+144). Median $log_2$ fold change at each timepoint is represented by the bold line.

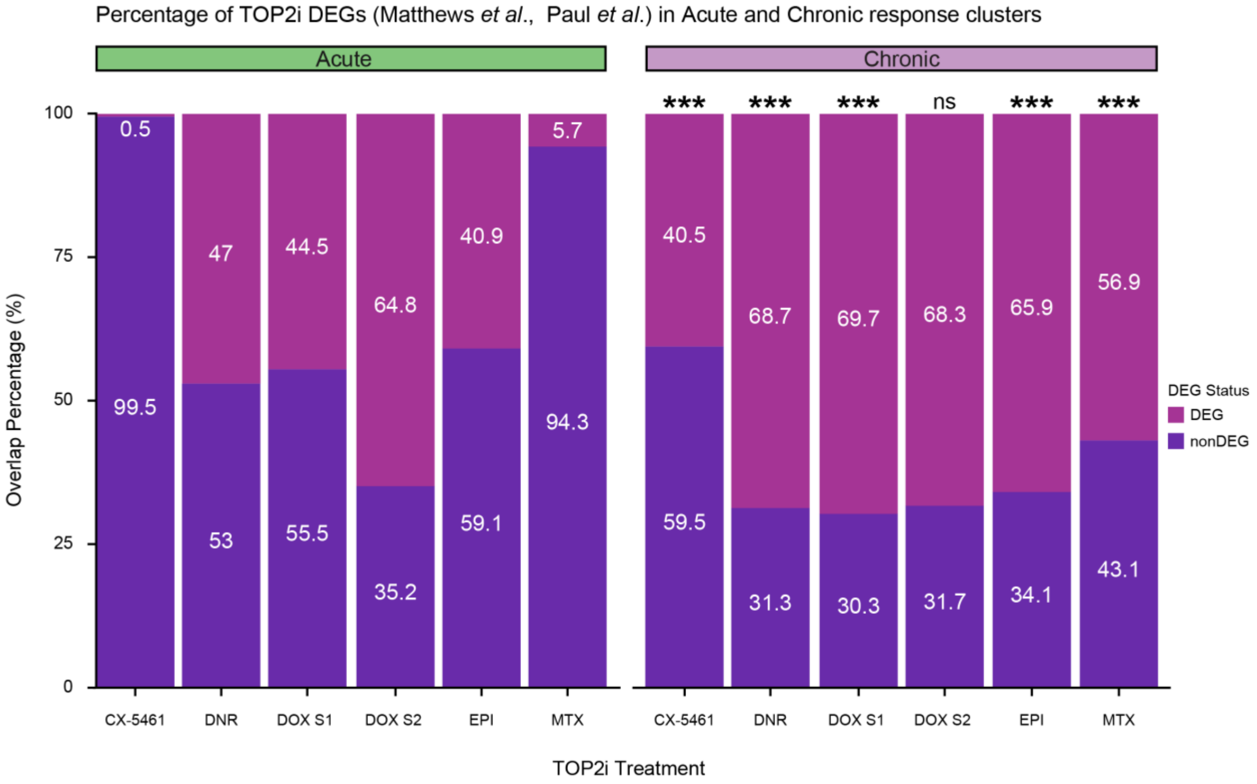


**Supplementary Figure 7: Chronic response genes are enriched in TOP2i DEGs compared to Acute response genes.** Percentage of Acute and Chronic response genes that overlap TOP2i DEGs from prior studies (Matthews *et al.*; Paul *et al.*): Daunorubicin (DNR), Epirubicin (EPI), Mitoxantrone (MTX) and DOX_S1 from Matthews *et al.* and CX-5461 and DOX_S2 from Paul *et al*. Asterisks represent TOP2i sets where the proportion in Chronic response cluster genes is greater than the Acute response cluster (***$P < 0.001$).

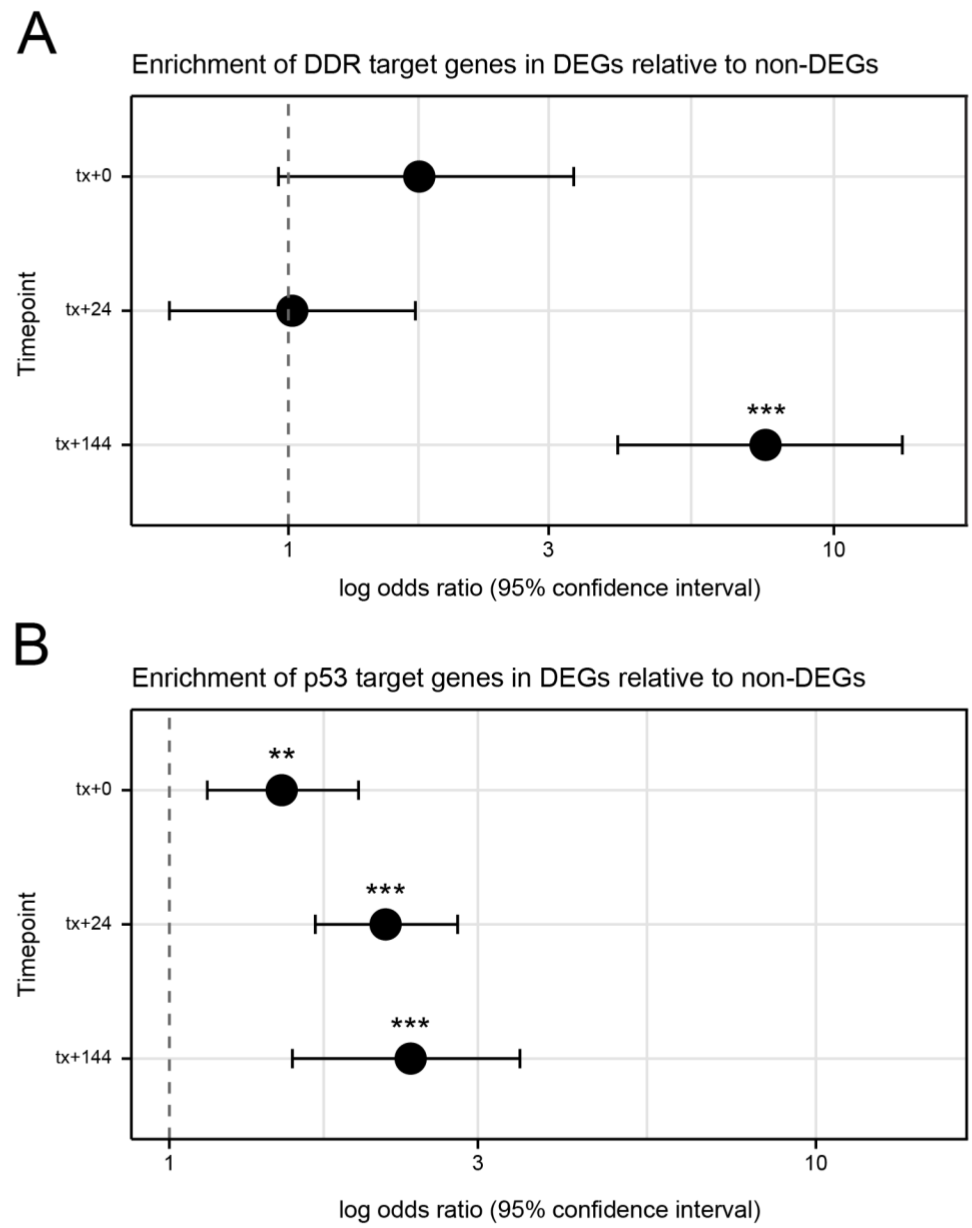


**Supplementary Figure 8: DDR and p53 target genes are enriched amongst late response genes. (A)** Enrichment of 65 DNA damage response (DDR) associated genes (Liberzon *et al.*) in DEGs compared to non-DEGs at each timepoint (tx+0, tx+24, and tx+144). Odds ratios from a Fisher's exact test are plotted together with 95% confidence intervals. Asterisk represents gene sets that are significantly enriched (**$P < 0.01$, ***$P < 0.001$). **(B)** Enrichment of p53 target genes (Fischer *et al.*) amongst DEGs compared to non-DEGs as described in (A).
